# Overcoming fixation and permeabilization challenges in flow cytometry by optical barcoding and multi-pass acquisition

**DOI:** 10.1101/2024.08.13.607771

**Authors:** Marissa D. Fahlberg, Sarah Forward, Emane Rose Assita, Michael Mazzola, Anna Kiem, Maris Handley, Seok-Hyun Yun, Sheldon J.J. Kwok

**Affiliations:** LASE Innovation Inc, Waltham, MA; Center for Regenerative Medicine, Massachusetts General Hospital, 185 Cambridge Street, Boston, MA 02114, USA; Harvard Stem Cell Institute, 7 Divinity Avenue, Cambridge, MA 02138, USA; Department of Stem Cell and Regenerative Biology, Harvard University, Cambridge, MA 02138, USA; Wellman Center for Photomedicine, Massachusetts General Hospital, Boston, MA

**Keywords:** optical barcoding, flow cytometry, laser particles, single cell, intracellular assays, fluorescent protein, phospho-flow, fixation and permeabilization

## Abstract

The fixation and permeabilization of cells are essential for labeling intracellular biomarkers in flow cytometry. However, these chemical treatments often alter fragile targets, such as cell surface and fluorescent proteins, and can destroy chemically-sensitive fluorescent labels. This reduces measurement accuracy and introduces compromises into sample workflows, leading to losses in data quality. Here, we demonstrate a novel multi-pass flow cytometry approach to address this long-standing problem. Our technique utilizes individual cell barcoding with laser particles, enabling sequential analysis of the same cells with single-cell resolution maintained. Chemically-fragile protein markers and their fluorochrome conjugates are measured prior to destructive sample processing and adjoined to subsequent measurements of intracellular markers after fixation and permeabilization. We demonstrate the effectiveness of our technique in accurately measuring intracellular fluorescent proteins and methanol-sensitive antigens and fluorophores, along with various surface and intracellular markers. This approach significantly enhances assay flexibility, enabling accurate and comprehensive cell analysis without the constraints of conventional one-time measurement flow cytometry. This innovation paves new avenues in flow cytometry for a wide range of applications in immuno-oncology, stem cell research, and cell biology.

## 1 Introduction

Over the past few decades, the scope of cell markers measured by flow cytometry has expanded from surface antigens to various intracellular proteins, such as cytokines and fluorescent reporter proteins, and intranuclear and genetic targets^1–10^. This expansion has enhanced the utility of flow cytometry across immunology and immuno-oncology^11–14^, stem cell research^15–17^, and cell biology^18–20^. For comprehensive phenotypic and functional analysis in these fields, it is essential to measure both surface and intracellular markers. However, cell processing for these measurements is challenging and can introduce significant measurement errors. For instance, fluorescent proteins (FPs) are commonly used to track gene uptake and expression, but fixation and permeabilization required to detect intracellular markers can cause physical loss and chemical alteration of intracellular FPs. Anti-GFP antibodies are used to mitigate this issue^21–23^, but they are often inadequate to recover the full signal from FPs. Similarly, when detecting phosphorylated proteins, methanol permeabilization damages the antigen epitopes of surface markers^24–27^,which are crucial for understanding cell signaling and immune response. These marker-destructive sample processing steps complicate assay design and prevent the optimal detection of markers.

Recently, we introduced multi-pass flow cytometry, which enables multiple measurements of the same cells using laser particles (LPs) as optical cell barcodes^28^. With each sequential flow measurement, different sets of markers are measured, and the data acquired are combined for each cell based on its unique barcode. We demonstrated a multi-pass workflow to acquire a 32-marker panel targeting surface markers with live cells^28^.

Here, we show that multi-pass flow cytometry offers effective solutions to long-standing difficulties associated with fixation, permeabilization, and methanol treatments in conventional flow cytometry. The key innovation enabled by cell barcoding is the ability to acquire sensitive and fragile markers first under optimal conditions. The sample is then processed for intracellular markers using methods which may be destructive to those measured in the first pass. The acquired data from the same cells through sequential flow cytometry are combined using the LP cell barcodes^28,29^. We demonstrate the compelling need for this approach and its effectiveness in three applications that require accurate measurement of methanol-sensitive epitopes, protein-based fluorophores, and fluorescent proteins in conjunction with harsh cell processing. Our method enables the detection of a wide range of previously incompatible marker types without compromise and risk of quantification errors.

## 2 Materials and Methods

### 2.1 Isolation of bone marrow cells from mice

All mice used in this study were maintained at Massachusetts General Hospital in a temperature- and humidity-controlled environment with a 12-hour light / 12-hour dark cycle and provided with food and water *ad libitum*. C57Bl/6-CAG-mRFP1-IRES-GFP mice, aged 8-12 weeks, were generated as previously described at the Harvard Genome Modification Facility^30^. The knock-in construct was modified from pR26CAG/GFP Dest (#74286, Addgene) by VectorBuilder to include a bicistronic fluorescent reporter encoding both mRFP and eGFP. Bone marrow cells were collected by crushing the tibias, femurs, hips, humeri, and spine of the mice. After collection, lineage cells were depleted with a lineage cell depletion kit (#130-090-858, Miltenyi Biotec) according to the manufacturer’s instructions. Enriched progenitors and hematopoietic stem cells were then utilized for further analysis. All experiments involving mice were conducted under the approval of the Institutional Animal Care and Use Committee at Massachusetts General Hospital (IACUC protocol #2016N000085).

### 2.2 MCF7 cell culture and intranuclear staining

MCF7 GFP- and GFP+ cell lines were sourced from ATCC (Manassas, VA) and GenTarget (San Diego, CA), respectively. These cells were cultured in MCF7 media (Minimal Essential Medium with 10% fetal bovine serum (FBS), 1% Penicillin/Streptomycin (P/S), 1% sodium pyruvate, and 1% non-essential amino acids (all (v/v)) in T75 flasks. Culturing was timed to allow passaging one day before experiment harvest. For detachment, 0.25% Trypsin ethylenediaminetetra-acetic acid (EDTA) was used and neutralized with MCF7 media. Cells were seeded into 12-well plates at ∼1.5 x 10^5^ cells/cm^2^, adjusting the volume to 2 mL per well with MCF7 media. Plates were incubated overnight at 37 °C with 5% CO_2_. After incubation, cells were harvested, counted, and allocated at ∼5 x 10^5^ cells per tube for each sample, reserving some GFP- and GFP+ cells as compensation controls.

For intranuclear staining, MCF7 cells were treated with a Foxp3/Transcription Factor Staining Buffer Set (eBioscience^TM^) following the manufacturer’s recommendations. In brief, samples were fixed and permeabilized with Foxp3 Fixation/Permeabilization working solution for 45 minutes at 4°C and washed twice with 1X Permeabilization Buffer prior to staining with Ki67-PE.

### 2.3 Multi-pass phospho-flow protocol

Cryopreserved human peripheral blood mononuclear cells (hPBMCs) were thawed and incubated in 0.1 mg/mL bovine pancreatic DNase I (STEMCELL) in Roswell Park Memorial Institute (RPMI) 1640 medium for 15 minutes at room temperature to mitigate cell clumping. Next, the hPBMCs were washed and resuspended in a 1:1000 dilution of LIVE/DEAD^TM^ Fixable Green Dead Cell Stain Kit (Invitrogen^TM^) in phosphate buffered saline (PBS), and incubated for 30 minutes in the dark at room temperature. The hPBMCs were then washed and resuspended in 325 µL of PBMC media (20% FBS (v/v), 1% P/S (v/v) in RPMI 1640). The cells were stimulated with 1X Cell Stimulation Cocktail (eBioscience^TM^), composed of PMA/Ionomycin, at 37°C for 15 minutes. After stimulation, the cells were immediately fixed by the addition of 200 µL of 4.2% formaldehyde (w/w) for 30 minutes at room temperature. Cells were washed in 2 mL of PBS, followed by another wash in 2 mL of wash buffer (10% FBS (v/v), 10 mM 2-[4-(2-hydroxyethyl)piperazin-1-yl]ethanesulfonic acid (HEPES) buffer, 2 mM EDTA, 1X poloxamer 188 non-ionic surfactant (Gibco^TM^) in PBS) at 600 g for 5 minutes each. Cells were then aliquoted into sample tubes for barcoding with LPs.

hPBMCs used in the phospho-flow protocol were barcoded after stimulation and fixation, prior to surface staining with antibodies. For barcoding, hPBMCs were stained with biotinylated antibodies against CD45 and β2-microglobulin (BioLegend) during a 15-minute incubation at 4°C. After washing, cells were resuspended in 1 mL of wash buffer. Streptavidin-coated LPs were added at a 10:1 LP:cell ratio. Samples were mixed using a HulaMixer™ (Invitrogen^TM^) at 4°C for 30 minutes, centrifuged at 600 g for 5 minutes, and the supernatant was discarded.

In 100 µL of wash buffer, samples were stained with an antibody panel targeting major cell populations (see Supplementary Table 1) in the dark at room temperature for 20 minutes. Following washing, samples were acquired and captured using a LASE multi-pass flow cytometer at a medium flow rate (30 µL/min). Captured samples were then fixed with 900 µL ice-cold methanol while vortexing, incubated on ice for 30 minutes, washed with PBS and wash buffer, and stained with p-ERK1/2 for 25 minutes in the dark at room temperature. After a final wash, cells were resuspended in 100 µL of wash buffer, and data from the second pass were acquired at a slow flow rate (10 µL/min) on the LASE multi-pass flow cytometer.

### 2.4 Multi-pass GFP / cell cycle protocol with MCF7 GFP+ and bone marrow cells

MCF7 cells were barcoded prior to staining with a 1:1000 dilution of LIVE/DEAD ^TM^ Fixable Near-IR Dead Cell Stain Kit (Invitrogen^TM^) in PBS for 30 minutes in the dark. To barcode, cells were harvested and resuspended in 100 µL of 0.1 mg/mL bovine pancreatic DNase I in RPMI 1640. A batch of polyethyleneimine (PEI) polymer-coated LPs were added to samples in 5 mL Eppendorf tubes once every 15 minutes, totaling to 4 additions, to achieve a final 10:1 LP:cell ratio. During this time, the sample tubes were mixing on an Eppendorf ThermoMixer® C at 300 rpm for 60 minutes at room temperature to facilitate the uniform, stochastic adhesion of LPs on the cell surface.

Mouse bone marrow cells were barcoded after antibody staining for lineage antibodies CD8A, CD3E, CD45R, GR1, CD11b, Ter119, and CD4 all conjugated to Alexa Fluor 700 (Supp. Table 1). To barcode, samples were resuspended in 100 µL of wash buffer containing a cocktail of biotinylated antibodies targeting progenitors and hematopoietic stem cells [H-2kb/H-2Db (Invitrogen^TM^), CD105 (eBioscience^TM^), CD150, CD45, and CD41 (BioLegend)] and incubated for 15 minutes at 4°C. After washing, samples were incubated in 100 µL of wash buffer containing 10 µg of purified streptavidin (BioLegend) for 25 minutes at 4°C. After a final wash, samples were resuspended in 1 mL of wash buffer. Biotin-coated LPs were added at a 10:1 LP:cell ratio. Samples were mixed at 5-minute intervals, alternating between centrifugation and thermo-mixing at 4°C for a total of 30 minutes. Finally, the cells were stained with viability dye.

All cells were acquired on a LASE multi-pass flow cytometer at a flow rate of 30 µL/min for the first pass, followed by cell capture. Captured samples were fixed and permeabilized in 250 µL of BD Fixation/Permeabilization solution (4.2% formaldehyde (w/w)) on ice for 20 minutes. The samples were washed and resuspended in 750 µL of 1X BD Perm/Wash^TM^ Buffer and centrifuged at 400 g for 5 minutes. Each pellet was resuspended in 500 µL of 1X BD Perm/Wash^TM^ Buffer, then stained with either Ki67-Alexa Fluor® 555 or Ki67-PE for 30 minutes in the dark on ice. Samples were washed with 500 µL of 1X BD Perm/Wash^TM^ Buffer and centrifuged at 400 g for 5 minutes. The pellets were resuspended in 500 µL of a 1:2500 dilution of 4′,6-diamidino-2-phenylindole (DAPI) in 1X BD Perm/Wash^TM^ Buffer and incubated in the dark on ice for 10 minutes. The samples were centrifuged at 400 g for 5 minutes and resuspended in 100 µL of wash buffer prior to a second acquisition at a flow rate of 10 µL/min.

### 2.5 Data analysis

Barcoded data were aligned using a proprietary matching algorithm and then exported as Flow Cytometry Standard (FCS) files for analysis using FlowJo version 10.10.0. Single-color compensation controls for all fluorescent antibodies, viability stains, DAPI, and fluorescent reporters were used for all experiments. Cytometer detector gains were set using detector setting incrementation to optimize the signal levels from single-color controls, ensuring they were the brightest in their respective channels for each run. Compensation controls were either as bright as or brighter than the corresponding samples. Compensation matrices from each pass were generated individually, automatically calculated with minimal manual modifications, and subsequently joined and applied to barcoded data for analysis. Compensation values for independently acquired fluorophore pairs were set to 0.

Data transformation and plotting were implemented using R software and the ‘tidyverse’ and ‘ggcyto’ packages^31–33^. A minimum of 100,000 events were collected per acquisition, with live single cells gated for subsequent analyses. In barcoded samples, only cells with a high statistical confidence of matching (< 5% error) were selected. For phospho-flow cytometry studies, gating was based on isotype controls and non-stimulated sample data. In experiments involving GFP and mRFP1, gating strategies were established using GFP-negative MCF7 cells and wild-type (non-reporter) mice as references. The calculation of fluorescent protein loss was conducted directly within FlowJo using the ratio of GFP and mRFP1 signals between the first and second passes.

## 3 Results

### 3.1 LP cell barcoding is compatible with harsh sample processing

We evaluated the effectiveness of multi-pass flow cytometry for measuring sets of distinctive markers that require competing sample processing and cell staining methods. Specifically, we investigated protocols requiring fixation and permeabilization, which are essential for staining DNA content, intranuclear proteins like transcription factors, phosphorylated proteins, and detecting fluorescent proteins within the cytoplasm. Fig. 1a shows a general workflow schematic. Cells are (1) stained with a panel of antibodies targeting surface markers or left untouched to detect fluorescent proteins, (2) barcoded with LPs, and (3) acquired through a LASE flow cytometer equipped with an LP barcode reader and a cell collector^28^. After the cells are captured, (4) they are fixed, permeabilized, and re-stained with intracellular markers, and (5) acquired in a second pass. Multi-pass data from LP-barcoded cells are matched, exported as an FCS file, and processed using conventional flow cytometry software.

**Figure 1.**
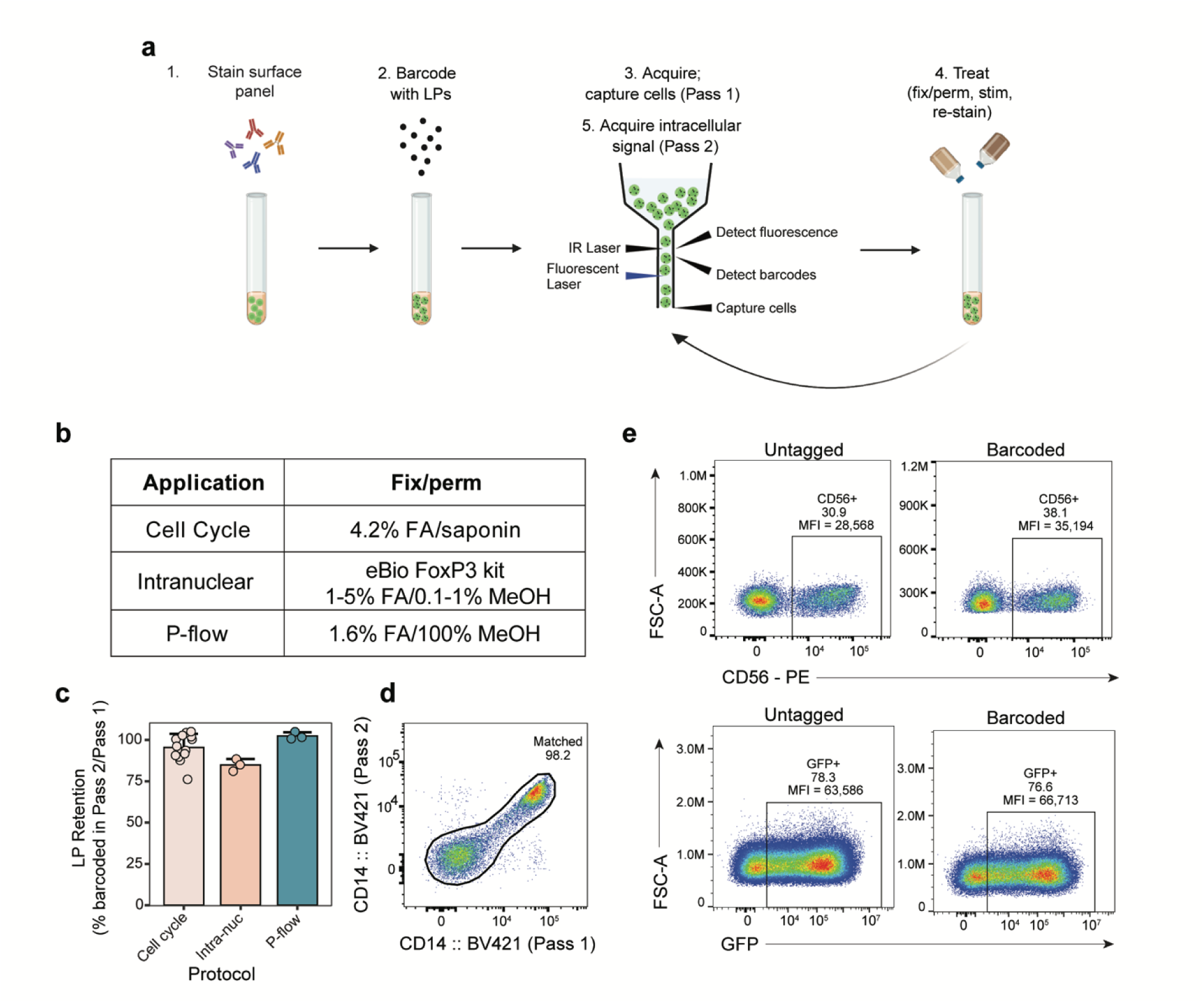
Robustness of LP cell barcoding across fix/perm protocols. (a) General schematic of the multi-pass workflow. Cells are typically stained for surface markers, barcoded with LPs, acquired through a LASE flow cytometer, captured, fix/perm treated, and re-analyzed. FA = formaldehyde. (b) Table showing three applications used for testing barcode stability and their respective fix/perm reagents. (c) Barcode retention in human PMBCs between passes in each application. Retention is calculated as percentage of cells barcoded in Pass 2 relative to the percentage of cells barcoded in Pass 1. (d) CD14 BV421 signals from LP-barcoded human PBMCs before (Pass 1) and after (Pass 2) the application of the phospho-flow (P-flow) protocol. Accurately matched cells with similar signal magnitudes fall along a diagonal axis. (e) Comparison of the fluorescent signals of a surface marker (CD56) and fluorescent protein (GFP) between untagged and matched cells from human PBMCs and MCF7 GFP+ cells, respectively.

We first examined the compatibility of LP barcoding across three commonly used, chemically harsh intracellular staining protocols on human PMBCs (Fig. 1b). The retention ratios of LP barcodes after various fixation and permeabilization (fix/perm) protocols were measured to be 95.4 ± 8.3% for the cell cycle workflow (4.2% formaldehyde (w/w)), 84.8 ± 3.7% for the intranuclear staining workflow (1-5% formaldehyde (w/w), 0.1-1% methanol (w/w)), and 102.4 ± 2.2% for the phospho-flow workflow (∼1.6% formaldehyde (w/w), 90% methanol (w/w)) (Fig. 1c). The barcode loss in the intranuclear staining workflow, which utilizes methanol as a strong permeabilizing agent instead of a saponin, a gentler permeabilizer used in standard fix/perm^34,35^, is likely attributed to partial detachment of LPs from the cell membrane. The accuracy of barcode matching was assessed by comparing the fluorescence intensities from anti-CD14 BV421 (Brilliant Violet 421^TM^) methanol-resistant antibodies stained on a human PBMC sample between the first and second measurements, that is, before and after the fix/perm phospho-flow protocol. A matching accuracy of over 98% was observed (Fig. 1d). Finally, the influence of LP-barcoding on surface marker and fluorescent protein expression was examined. The frequency and median fluorescent intensity (MFI) of surface marker and GFP+ events were within the typical coefficients of variation (CVs) of the assay and cytometry for both LP-barcoded and untagged cells (Fig. 1e).

### 3.2 Measurement of protein fluorophores before methanol treatment

We developed a two-pass phospho-flow protocol that circumvents the harsh effects of methanol on protein-based fluorophores (Fig. 2a) and tested its effectiveness for analyzing PMA/ionomycin-stimulated phosphorylation of ERK1/2 as a model system. Cryopreserved human PBMCs were (1) thawed and stained with viability dye, followed by (2) stimulation with PMA/Ionomycin for 15 minutes and (3) immediate fixation to preserve the integrity of the phosphorylated protein states. PBMCs were then (4) barcoded with LPs and (5) stained with antibodies conjugated to protein-based fluorophores targeting broad immune cell populations (CD3 PE-Cy5, CD20 APC-Cy7, CD14 BV421, CD56 PE, and HLA-DR PE-Dazzle594). Cells were then (6) acquired through an LP-enabled flow cytometer, captured, (7) permeabilized with 90% ice-cold methanol (v/v), (8) stained intracellularly for p-ERK1/2, and (9) re-acquired in a second pass.

**Figure 2.**
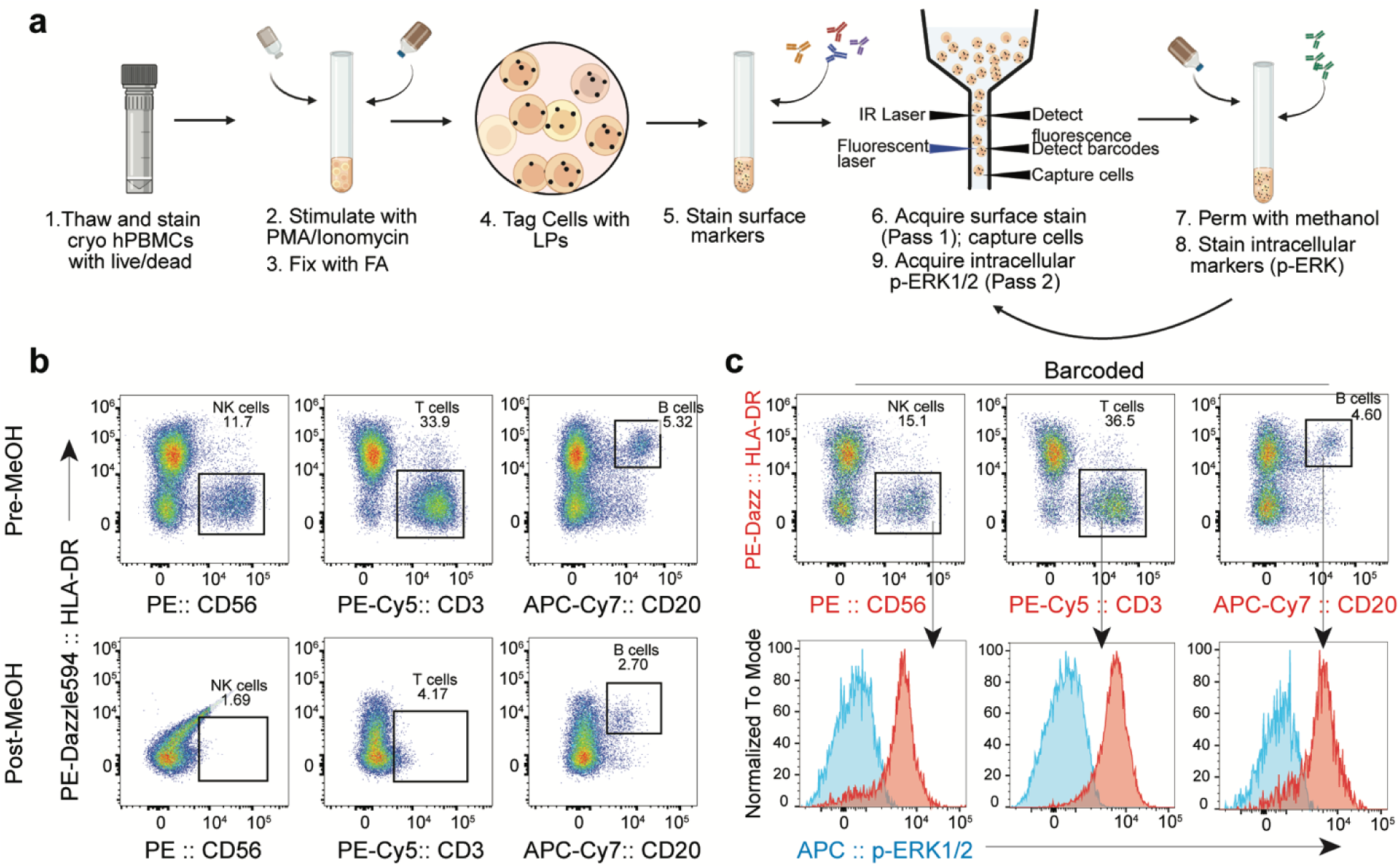
Full expression of fluorescence signal measured prior to methanol permeabilization. (a) Schematic of the multi-pass phospho-flow protocol for human PBMCs stimulated with PMA/Ionomycin. (b) Direct comparison of fluorescence from unmatched, barcoded cells before (top) and after (bottom) methanol permeabilization, showing degradation of protein-based fluorophore signals by the MeOH (methanol) treatment. (c) (top) Intact surface marker data acquired in the first-pass flow cytometry and gating of major cell populations; (bottom) Downstream p-ERK1/2 analysis from intracellular data collected during the second pass post fix/perm, gated off barcode-matched, pre fix/perm surface markers.

We found that fluorescence from PE- and APC-based fluorophores were significantly reduced or destroyed after methanol permeabilization, consistent with literature reports^26,36^. This loss of sensitivity resulted in partial (CD20 APC-Cy7) or complete (CD56 PE and CD3 PE-Cy5) inability to identify major cell populations (Fig. 2b). In contrast, measuring these markers in the first pass before methanol permeabilization allowed us to identify these markers without the detrimental effects of methanol. In the second pass, the intracellular p-ERK1/2 marker was measured and analyzed with respect to different cell populations identified by the data measured in the first pass prior to methanol permeabilization (Fig. 2c). The population percentages and MFI of all fluorophores between barcoded and untagged data were within typical CVs of the assay and cytometer (Figs. 2b-c).

### 3.3 Measurement of sensitive epitopes before methanol permeabilization

Traditionally, phospho-flow studies have been restricted to using small-molecule fluorophores that are resistant to methanol. However, significant issues arise when measuring surface markers that are sensitive to methanol treatment^26,37–42^. For example, the crucial antigen CD19 is highly susceptible to methanol denaturation. Two common protocols have been developed to partially mitigate this issue: CD19 is stained prior to methanol permeabilization with a methanol-resistant fluorophore (SOP1)^24^ or CD19 is stained after methanol permeabilization with any fluorophore (SOP2)^43^, as illustrated in Fig. 3a. However, both approaches are suboptimal and significantly compromise signal quality^43^. Our experiments revealed a large (25%) failure in detecting CD19+ cells (Fig. 3b) due to the denaturation of the CD19 antigen in these protocols.

**Figure 3.**
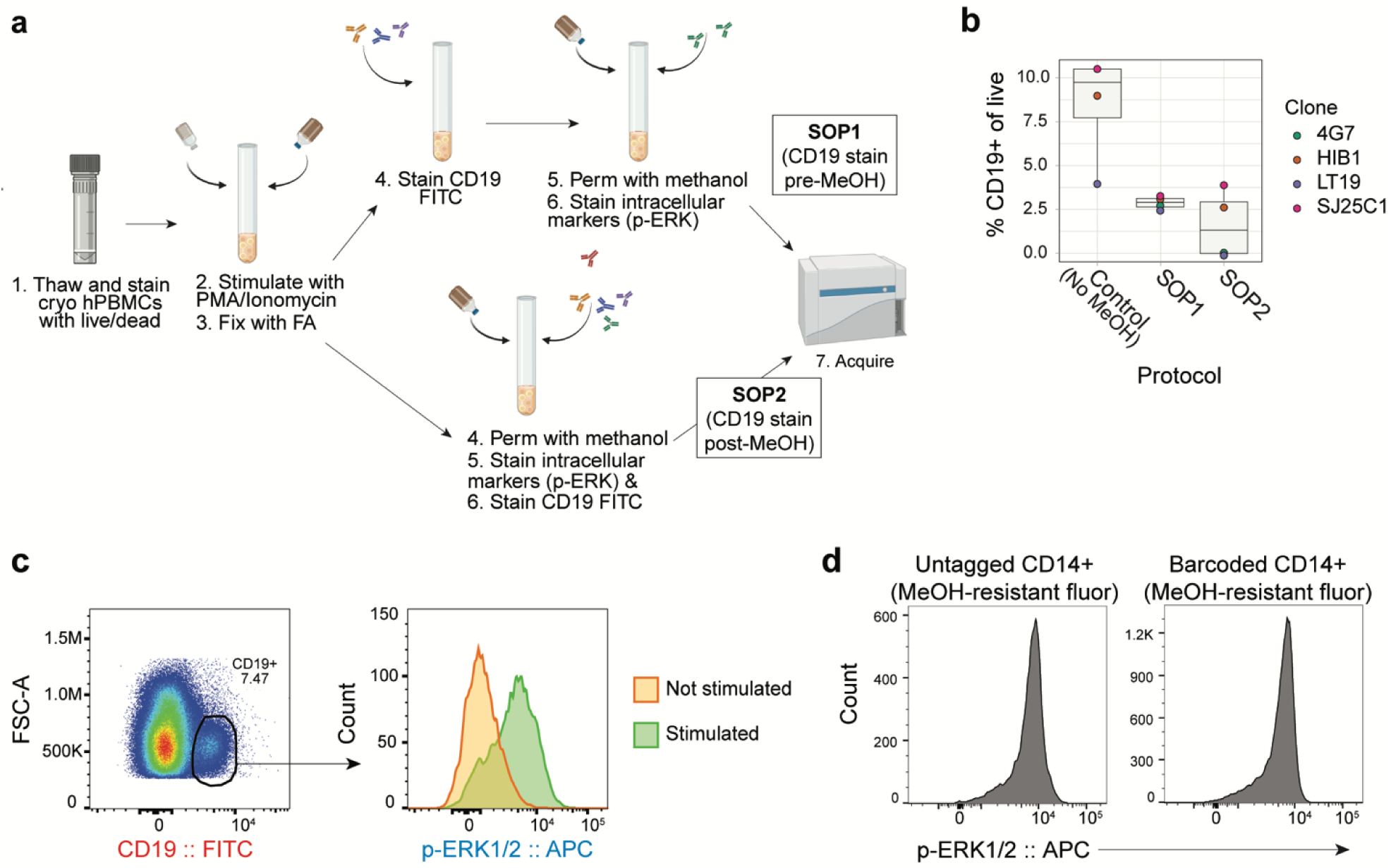
Acquisition of CD19 antigen data prior to its destruction by methanol permeabilization. (a) Schematic of two conventional protocols (SOP1 and SOP2) used to detect CD19 expression after fixation and methanol permeabilization. (b) The percentage of CD19+ cells out of total live cells using the conventional methods. (c) (Left) CD19 data acquired in the first pass (red text); (Right) p-ERK1/2 expression acquired during the second pass (blue text) for CD19+ cells identified by the intact CD19 data in the first pass. (d) Comparison of intracellular p-ERK1/2 expression on CD14+ cells stained with methanol-resistant BV421 between untagged and barcoded samples.

To solve this problem, we developed a two-pass protocol, which measures CD19 in the first flow pass prior to methanol permeabilization and then intracellular signals in the second pass. The combined data provide the p-ERK1/2 expression levels of all CD19+ cells (Fig. 3c). We found no difference in p-ERK1/2 expression between LP-tagged and untagged CD14+ monocytes (Fig. 3d).

### 3.4 Measurement of fluorescent proteins before fix/perm

Genetically encoded FPs are commonly used to analyze gene insertion and expression, but their stability is vulnerable to fix/perm methods. This poses challenges in accurately quantifying FP expression alongside intracellular marker signals. We used multi-pass flow cytometry to capture complete FP signals before fix/perm and intracellular marker signals after fix/perm, utilizing GFP- expressing MCF7 breast cancer cell lines and GFP/mRFP1 co-expressing murine hematopoietic stem and progenitor cells (HSPCs). Cells were stained with viability dye and/or surface antibody-fluorophores followed by barcoding with LPs (Fig. 4a). After the first pass, the collected cells underwent fix/perm using various reagents and protocols, followed by staining with an intracellular cell cycle dye and the second pass.

**Figure 4.**
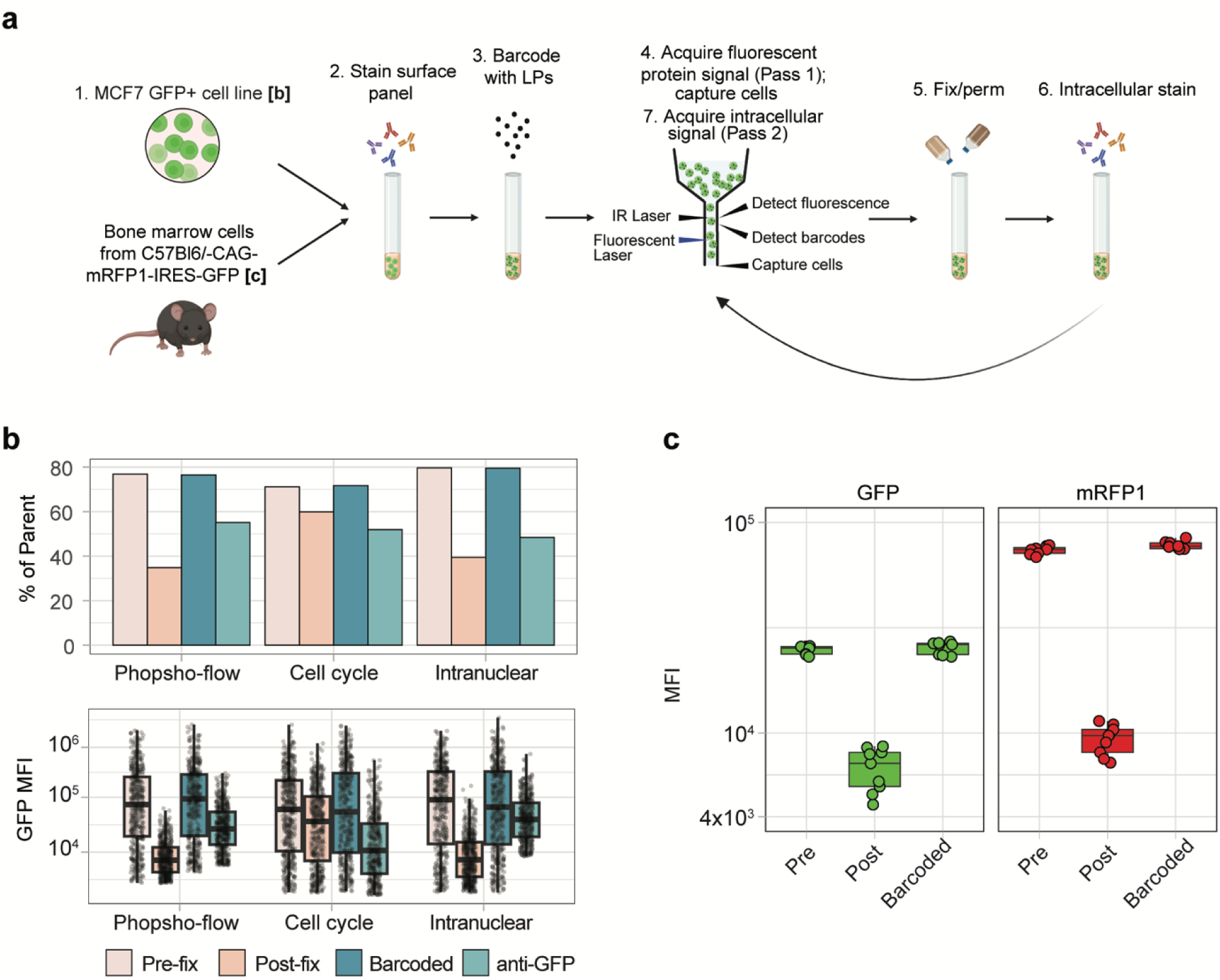
Measurement of intact fluorescent protein expression using multi-pass flow cytometry. (a) Schematic of the multi-pass workflow for acquisition of fluorescent protein expression on live cells prior to fix/perm and intracellular stain. (b) Percentage and MFI of GFP from MCF7 GFP+ cells detected pre-fix/perm, post-fix/perm, using two-pass analysis protocol, and using anti-GFP APC following three different fix/perm protocols. (c) MFI of GFP/mRFP1 co-expressing bone marrow cells measured before cell cycle fix/perm without barcoding, after fix/perm without barcoding, and with the two-pass protocol with barcoding. Data were collected in triplicate from three mouse donors, with each point representing one replicate.

For MCF7 GFP+ cells, we observed a 50% loss of GFP+ events after employing the intranuclear and phospho-flow protocols. Additionally, more than 10% of GFP+ events were lost during cell cycle processing (Fig. 4b). These protocols also similarly decreased MFIs. One method of retaining GFP expression is through an anti-GFP antibody; however, this approach only partially restored the frequencies to 60-72% of the originals and GFP MFI was significantly reduced (Fig. 4b).

Similarly, when applying the cell cycle fix/perm protocol to bone marrow stained to demark immunophenotypic HSPCs, both GFP and mRFP1 signals experienced a significant decrease post fix/perm. Specifically, the MFI of GFP and mRFP1 after fix/perm dropped to 27% and 9% of their original levels, respectively, and differential signal loss between the two FPs was observed. In contrast, measuring FPs prior to fix/perm treatments in our multi-pass workflow fully restored the FP signals (Fig. 4c). Therefore, our method allows quantitation of expression in combination with intra- and extracellular stains that require fixation/permeabilization. Therefore, our method allows quantitation of multiple FP expression in combination with intra- and extracellular stains that require fixation/permeabilization.

### 3.5 Multi-pass analysis reveals differential fluorescent protein signal loss

Using our model system, we investigated the loss of FP signal due to fix/perm treatments. First, we verified that LP tagging does not alter the expression of intracellular cell cycle signals in MCF7 cells (Fig. 5a). Next, using our single-cell barcoding workflow we compared GFP expression in MCF7 cells before and after fix/perm. This comparison confirmed the previously noted losses and identified specific cells that exhibited undetectable levels of GFP in the second pass (Fig. 5b). By computing the ratio of each cell’s GFP fluorescence intensity in the second pass to the fluorescence intensity of GFP in the first pass, we found that that nearly 80% of cells underwent a 50% or greater loss of GFP (Fig. 5c).

**Figure 5.**
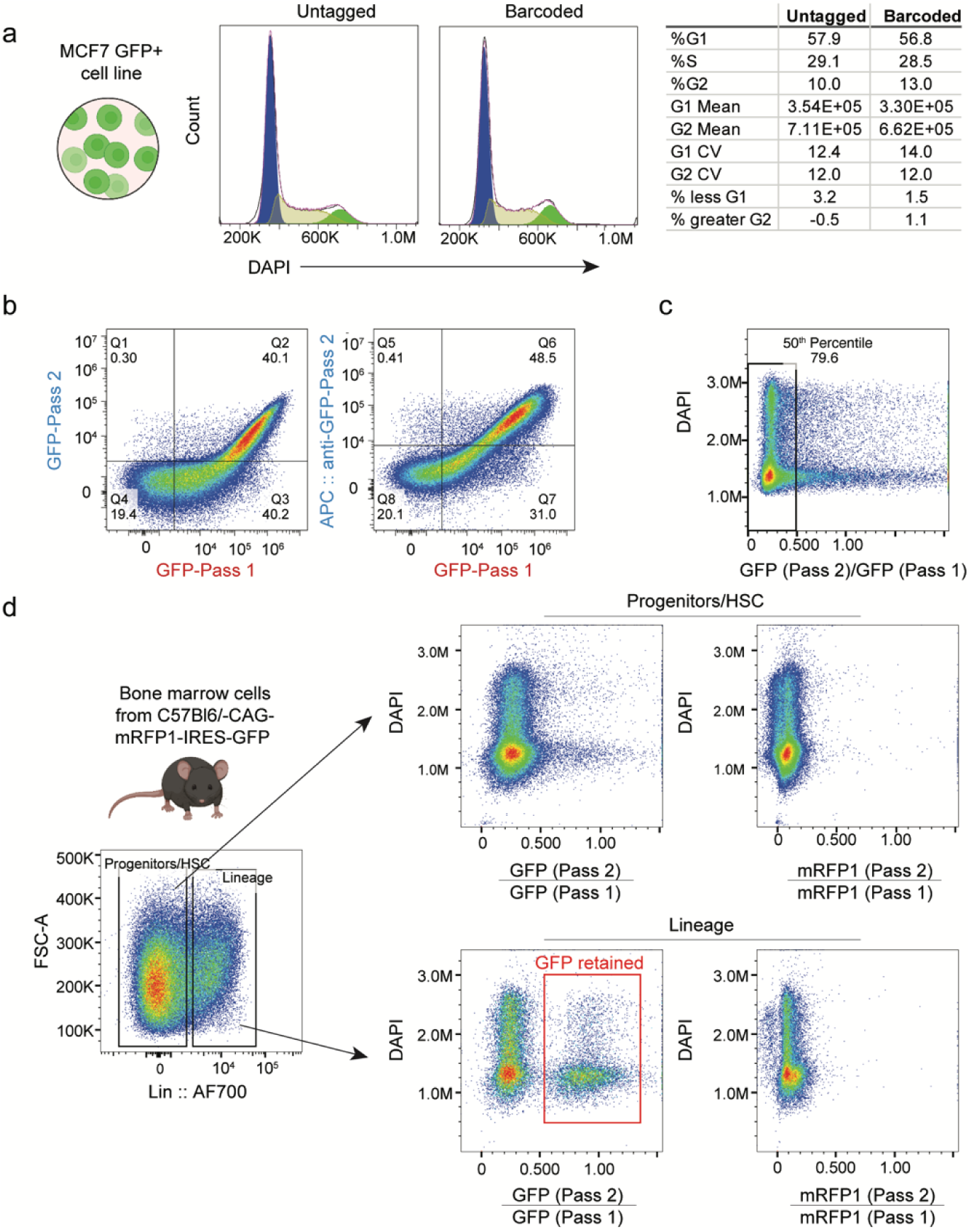
Fluorescent protein cell loss differs by cell type and cell cycle stage. (a) Representative histograms comparing DAPI expression on untagged and LP-tagged MCF7 GFP+ cells. (b) Direct comparison of GFP expression from MCF7 cells before and after fix/perm (intranuclear protocol) and relative to the anti-GFP antibody stained after fix/perm. Red text highlights data collected in the first pass, and blue text represents data collected in the second pass. (c) Representative dot plot showing the GFP signal change between the first and second passes. Most cells significantly lost GFP signals in the second pass after fix/perm. (d) (Left) marker gating data and the loss of GFP and mRFP1 across (top) hematopoietic progenitors and stem cells (HSC) and (bottom) lineage+ populations before and after fix/perm.

Noting the differential loss in GFP and mRFP1 expression in murine HSPCs, we investigated whether FP loss was dependent on cell type. Significant reduction in GFP and mRFP1 intensities were observed among both HSPCs (Fig. 5d). However, we observed a distinct cell population among lineage-positive cells that maintained a similar amount of GFP expression across passes, which was not observed with mRFP1. These results highlight a potential source of bias if FP signals are only quantified post fix/perm.

## 4 Discussion

In flow cytometry, detecting each marker type often requires specific sample processing methods that may interfere with the detection of other markers. This issue frequently arises in immuneoncology and stem cell biology research. For example, detecting phosphorylated signaling proteins and fragile surface markers on the same cells is essential for studying cell division and survival in cancer, but methanol permeabilization required for phospho-flow can degrade surface markers needed for phenotyping^3^. To address this problem, researchers often optimize fixative concentrations and incubation times to enable simultaneous detection of both types of markers. However, this approach inevitably compromises signal quality for each marker type, ultimately reducing assay sensitivity^21,24,25^. Another strategy involves designing panels using only methanol-resistant fluorochromes and epitopes, but this complicates and limits panel design and may necessitate the exclusion of important markers^27^. Alternatively, pre-sorting cells based on surface marker expression, followed by intracellular staining and re-acquisition, is sometimes used. However, this method is time-intensive and costly, and impractical for sorting many cell types at once, leading to the loss of single-cell data collected during sorting.

We have presented a novel multi-pass approach that resolves this previous dilemma by enabling separate flow measurements of different markers. Optical barcoding using stable laser particles allows repeated measurements of the same cells. Instead of measuring every marker of interest simultaneously, cells can be acquired before and after each cell processing step, with the resulting data from each pass integrated for each cell via barcoding. This allows workflows optimized for the detection of each marker type to be used without compromise.

We demonstrated several applications of our approach. First, we showed surface phenotyping of stimulated human PBMCs combined with the detection of phosphorylated protein p-ERK1/2 by acquiring cells before and after methanol permeabilization. Second, we showed detection of FP expression from MCF7 and mouse bone marrow cells combined with intracellular cell cycle staining by acquiring cells before and after fixation and permeabilization. In both cases, samples processed through the multi-pass workflow produced a single FCS file that integrated single-cell data from fix/perm-incompatible signals obtained during the first pass (FP, sensitive fluorochromes, and fragile epitopes) with cell cycle or phosphorylation data acquired in the second pass. For all workflows, we established the stability of barcoding and confirmed that laser particle tagging did not influence the detection or quality of FP or intracellular data. The loss of GFP and mRFP1 with fix/perm appears to be dependent on FP type, cell type, and cell cycle stage.

Our multi-pass flow cytometry approach allows for flexible and simplified panel design through unencumbered fluorochrome and epitope choice, and significant savings on resources spent optimizing assay-specific parameters, including antibody clones, fluorochromes, buffers, reagent concentrations, and workflows. We anticipate this method will facilitate unprecedented cellular analysis from phenotype to state to function through optimized detection of different marker types on the same cells.

## Author Contributions

S.J.J.K., M.D.F, S.F., M.M, and M.H designed the study. S.F., E.R.A., M.M., and A.K. performed flow cytometry experiments. S.J.J.K., M.D.F, S.F., M.M., and E.R.A. analyzed and interpreted data.

S.J.J.K. and M.D.F. prepared the manuscript with input from all authors.

