## Supplementary Table 1 for "Overcoming fixation and permeabilization challenges in flow cytometry by optical barcoding and multi-pass acquisition"

| **Antigen/Target** | **Fluorophore** | **Catalog no.** | **Clone** | **Supplier** |
| --- | --- | --- | --- | --- |
| DNA | DAPI | 422801 | - | BioLegend |
| CD44 | Brilliant Violet 421^TM^ | 338809 | BJI8 | BioLegend |
| HLA-DR | PE/Dazzle^TM^ 594 | 307653 | L243 | BioLegend |
| CD14 | Brilliant Violet 421^TM^ | 301829 | M5E2 | BioLegend |
| CD56 (NCAM) | PE | 318306 | HCD56 | BioLegend |
| CD45 | PerCP | 304025 | HI30 | BioLegend |
| GFP | APC | 338010 | FM264G | BioLegend |
| ERK1/2 Phospho (Thr202/Tyr204) | APC | 369521 | 6B8B69 | BioLegend |
| CD3 | PE/Cy^TM^5 | 300310 | HIT3a | BioLegend |
| CD19 | FITC | 363007 | SJ25C1 | BioLegend |
| CD19 | FITC | 392507 | 4G7 | BioLegend |
| CD19 | FITC | 302205 | HIB19 | BioLegend |
| CD19 | FITC | 130-113-730 | LT19 | Miltenyi Biotec |
| Ki-67 | PE | 652404 | 16A8 | BioLegend |
| CD8A | Biotin | 553029 | 53-6.7 | BD Biosciences |
| CD3E | Biotin | 553060 | 145-2C11 | BD Biosciences |
| CD45R | Biotin | 553086 | RA3-6B2 | BD Biosciences |
| GR1 | Biotin | 553125 | RB6-8C5 | BD Biosciences |
| CD11b | Biotin | 553309 | M1/70 | BD Biosciences |
| Ter119 | Biotin | 553672 | Ter-119 | BD Biosciences |
| CD4 | Biotin | 553728 | GK1.5 | BD Biosciences |
| Streptavidin | AF700 | S21383 | - | Invitrogen |
| H-2kb/H-2Db | Biotin | PIMA517998 | 5041.16.1 | Invitrogen |
| CD105 | Biotin | 13-1051-85 | MJ7/18 | eBioscience |
| CD150 | Biotin | 115908 | TC15-12F12.2 | BioLegend |
| CD45 | Biotin | 103104 | 30-F11 | BioLegend |
| CD41 | Biotin | 133930 | MWReg30 | BioLegend |

**Supplementary Table 1.** Details of reagents used in this study.
